# Spatiotemporal spinal integration of descending and spinal volleys in spinal motor circuits revealed by compound motor evoked potentials

**DOI:** 10.64898/2026.06.02.729725

**Authors:** Yousuke Tanaka, Atsushi Sasaki, Nadaka Hakariya, Hiroki Arakawa, Yume Mashiki, Ryo Aoki, Youhei Masugi, Dimitry G. Sayenko, Kimitaka Nakazawa

## Abstract

Descending corticospinal and afferent pathways underlying spinally evoked motor potential both contribute to motor output, yet how their interaction at the spinal and peripheral levels is organized spatially within a muscle remains unclear. This study investigated the spatiotemporal characteristics of descending modulation of spinally evoked motor potentials by combining subthreshold transcranial magnetic stimulation (TMS) with transcutaneous spinal cord stimulation (tSCS). In Experiment 1, spinally evoked motor potentials were recorded from multiple lower-limb muscles at various interstimulus intervals (ISIs) defined relative to central conduction time (CCT). Subthreshold TMS facilitated spinally evoked motor potentials from CCT onward across all recorded muscles, with additional bilateral facilitation observed at longer ISIs. In Experiment 2, high-density surface electromyography (HDsEMG) revealed distinct intramuscular activation patterns in the tibialis anterior. The center of gravity (CoG) of TMS-induced motor evoked potentials was located more proximally than that of spinally evoked motor potentials. Notably, the CoG of facilitation maps was shifted further proximally than that of both single-stimulus responses. These findings suggest that descending and afferent inputs preferentially recruit partially distinct motoneuron pools within the same muscle. The proximal bias of facilitation indicates recruitment of additional motoneurons rather than uniform amplification of existing activity. Together, these results demonstrate that the interaction between descending and afferent inputs is both timing-dependent and spatially non-uniform, providing new insight into sensorimotor integration in the human lower limb.

## Introduction

Human motor output is shaped by multiple neural mechanisms, among which modulation of descending voluntary drive from the primary motor cortex and regulation of spinal interneuron and motoneuron excitability are two major contributors (Kandel et al. 2000; Knikou and Taglianetti 2006; Pierrot-Deseilligny and Burke 2005). Both mechanisms contribute to gait (Capaday and Stein 1986; Dietz et al. 1990; Yang and Whelan 1993) and postural control (Koceja et al. 1995; Tokuno et al. 2007, 2008; Zehr and Stein 1999), and their interaction shapes motor output in a context-dependent manner. Since voluntary commands reach spinal motoneurons largely via corticospinal projections (Glover and Baker 2022; Kandel et al. 2000; Usuda et al. 2022), a central question is how corticospinal drive is integrated with spinal motoneuronal and interneuronal circuits in a spatiotemporally structured manner. To examine corticospinal influences on spinal network, previous studies have primarily combined transcranial magnetic stimulation (TMS) over the primary motor cortex with the soleus H-reflex, thereby characterizing the effects of descending input on reflex excitability primarily in a single muscle (Andrews et al. 2020). These studies have demonstrated that a conditioning TMS pulse facilitates H-reflex amplitude in a timing-dependent manner, and that similar facilitation can occur even when TMS is delivered below motor threshold (Andrews et al. 2020; Nielsen and Petersen 1995; Rossi et al. 2025). Together, these findings suggest that TMS-induced descending volleys transiently increase motoneuron excitability, such that subsequent afferent input produces a larger reflex response.

However, single-muscle TMS-H-reflex studies do not capture the multi-muscle, multi-segmental organization of spinal circuits. Single-muscle H-reflex paradigms therefore provide a limited perspective on how descending input modulates reflex pathways at the system level. Moreover, while previous studies have largely defined the temporal dependence of this interaction (Andrews et al. 2020; Nielsen and Petersen 1995; Rossi et al. 2025), its spatial organization remains unclear, particularly with respect to how modulation is distributed across spinal segments and expressed within individual muscles.

To address these issues, we applied transcutaneous spinal cord stimulation (tSCS) over the thoracolumbar region to elicit spinally evoked motor potentials concurrently in multiple lower-limb muscles (Courtine et al. 2007; Danner et al. 2016; Kaneko et al. 2021; Saito et al. 2019; Sasaki et al. 2020; Sayenko et al. 2015a). Spinally evoked motor potentials are modulated in a timing-dependent manner when combined with TMS (Roy et al. 2014; Sayenko et al. 2018), and such modulation may not be restricted to the target muscle but also extends to non-target and contralateral muscles, reflecting that descending influences on spinal reflex pathways are not confined to a single muscle (Roy et al. 2014; Sayenko et al. 2018; Tazoe and Perez 2014; Ziemann et al. 1999). Therefore, evaluating modulation across multiple muscles may provide important insight into how descending inputs are distributed and integrated within bilateral and multi-muscle spinal motor networks involved in locomotion and postural control.

However, intermuscular modulation patterns alone provide limited mechanistic insight, as compound motor evoked potential (MEP) amplitude reflects response magnitude but do not distinguish whether TMS-induced modulation of spinal responses arises from a uniform increase in the excitability of existing motoneurons and interneurons or from the recruitment of additional motor units. We therefore employed high-density surface electromyography (HDsEMG) in the tibialis anterior (TA), the target muscle for TMS, to examine the intramuscular spatial distribution during single (TMS-only and tSCS-only) and combined stimulation. Because the TA receives stronger corticospinal projections than the thigh muscles and soleus (Bawa et al. 2002; Brouwer and Ashby 1992; Eisner-Janowicz et al. 2023), TMS-induced modulation of spinally evoked motor potentials may be more readily observed in this muscle, making it a suitable model for investigating the underlying mechanisms of the modulation. However, since the TA is a muscle in which H-reflexes are difficult to elicit, the modulatory effects of TMS on spinally evoked motor potentials in this muscle have not been sufficiently investigated. Previous studies have shown that differences in neural input to motoneurons can shift the center of intramuscular activity, and such shifts are indicative of changes in motor unit recruitment (Elswijk et al. 2008; Falla and Gallina 2020; Farina et al. 2008). Therefore, we compared the spatial distributions of TMS-induced MEPs and spinally evoked motor potentials within the TA to identify the intramuscular regions in which facilitation emerged during combined stimulation. This approach allowed us to determine whether the interaction between descending and afferent inputs is expressed uniformly across the muscle or preferentially in specific intramuscular regions. Because descending and afferent inputs may converge non-uniformly within the motoneuron pool (Morita et al. 1999), modulation of the compound MEP during combined stimulation may reflect the preferential weighting of partially overlapping motoneuron populations by descending and afferent inputs. Given that the composition of motoneuron pools varies within the TA muscle (Okuma et al. 2013; Weinman et al. 2026), and that the recruitment of distinct motoneuron pools can produce different intramuscular activation patterns (Falla and Gallina 2020; Farina et al. 2008), the modulation of the compound MEP during combined stimulation may likewise be spatially non-uniform within the muscle. The spatial distribution of the compound MEP within a muscle may therefore provide insight into how descending and afferent inputs are integrated at the spinal level.

The aims of the present study were: (1) to investigate the timing-dependent effects of TMS on spinally evoked motor potentials across multiple lower-limb muscles bilaterally, and to characterize the intermuscular distribution of reflex modulation (*Exp. 1*); and (2) to characterize the intramuscular spatial distribution of the modulated responses using HDsEMG (*Exp. 2*). We hypothesized that: (1) spinally evoked motor potentials would be modulated in a timing-dependent and most pronounced in the target muscle (i.e., TA), with possible effects in non-target muscles; and (2) modulation during combined stimulation would be spatially non-uniform within the target muscle, indicating that the interaction between descending and afferent inputs is not expressed uniformly at the spinal level.

## Methods

### Participants

Ten healthy male participants [age: 25.0 ± 4.0 yr, height: 1.72 ± 0.07 m, body mass: 66.6 ± 8.6 kg (mean ± SD)] participated in both *Exp. 1* and *Exp. 2*. None of the participants had a history of neurological or musculoskeletal impairments. All participants provided written informed consent in accordance with the Declaration of Helsinki. The experimental procedures were approved by the institutional ethics committee of The University of Tokyo (533-9).

### General settings

Participants remained relaxed in the supine position throughout the experiments to minimize the influence of posture and antigravity muscle activity on spinally evoked motor potentials responsiveness (Danner et al. 2016; Koceja et al. 1995) (Figure 1). Both legs were stabilized using ankle-foot orthoses and straps to prevent movements during the experiments.

**Figure 1.**
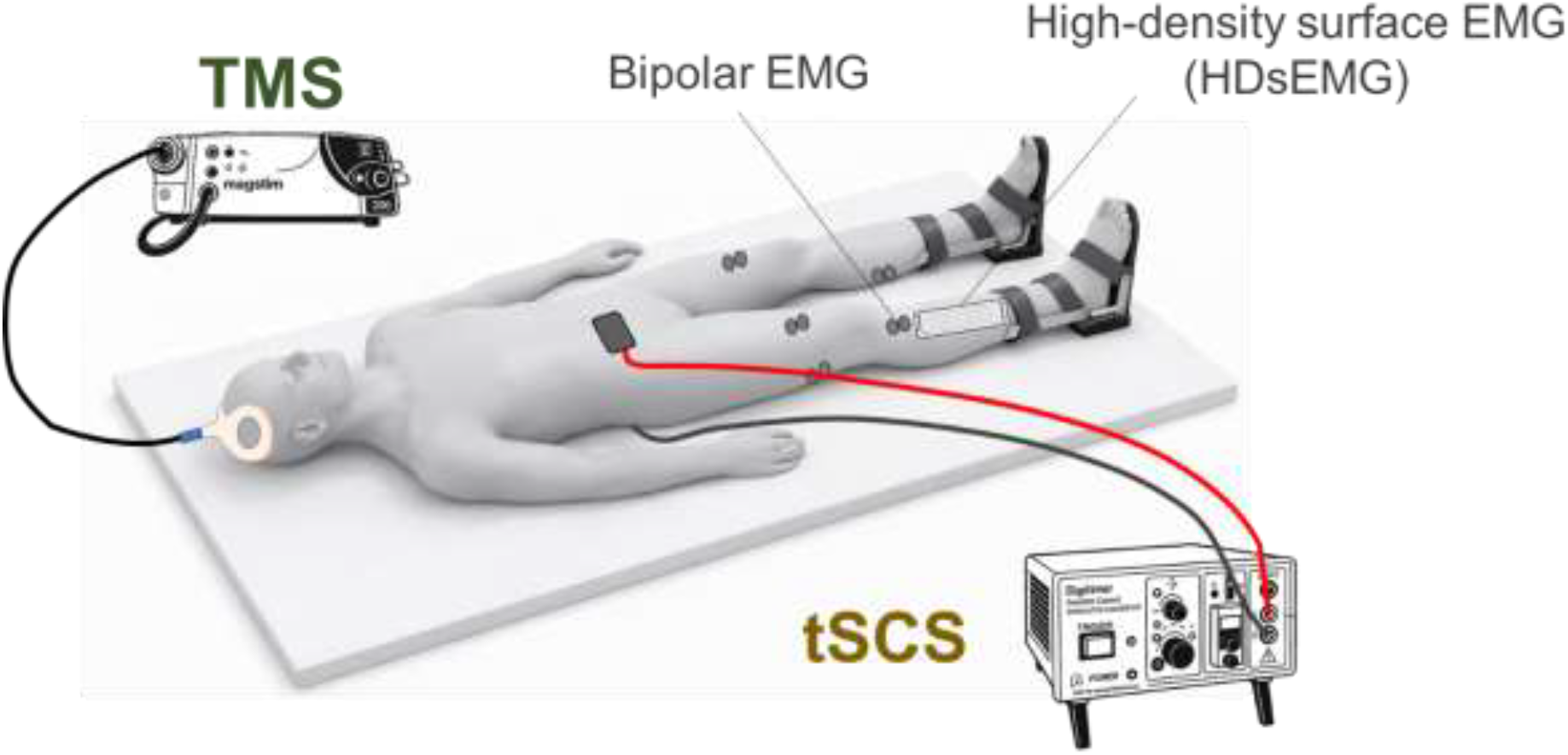
Experimental setup. Participants were positioned supine during the experiment. The figure illustrates the transcranial magnetic stimulation (TMS), the transcutaneous spinal cord stimulation (tSCS) electrodes, bipolar EMG electrodes, and high-density surface EMG (HDsEMG). Participants remained relaxed throughout the experiment.

### EMG settings

Bipolar Ag/AgCl surface electrodes (Vitrode F-150S; Nihon Kohden, Tokyo, Japan) were placed over the muscles according to SENIAM recommendations (Hermens et al. 2000) after the skin had been cleaned with alcohol. The interelectrode distance was approximately 1 cm. A ground electrode was placed over the right lateral malleolus. Bipolar EMG signals were amplified (×1,000) with a multichannel EMG amplifier (MEG-6108; Nihon Kohden, Tokyo, Japan), digitized at 10 kHz with an analog-to-digital (A/D) converter (PowerLab/16SP; AD Instruments, Castle Hill, Australia), and stored on a computer for offline analysis. Bipolar EMG was recorded bilaterally from the TA, medial gastrocnemius (MG), rectus femoris (RF), and biceps femoris (BF) to assess the net effects of TMS conditioning on spinally evoked motor potentials.

HDsEMG was recorded from the right TA using a 64-channel grid electrode (GR08MM1305; OT Bioelettronica, Torino, Italy) to evaluate spatial patterns of intramuscular activation. The grid comprised 64 electrodes arranged in a 13 × 5 grid configuration, with one corner position left empty by design (electrode diameter: 2 mm; interelectrode distance: 8 mm). Signals were sampled at 2,000 Hz. To ensure consistent electrode placement across participants, the grid was positioned according to SENIAM recommendations (Hermens et al. 2000): (i) the sixth row of the HDsEMG grid was aligned with a point located one-third of the distance from the fibular head to the medial malleolus; (ii) the grid was oriented along the longitudinal fiber direction of the TA; and (iii) the medial edge of the grid was aligned with the lateral border of the tibia. Because of the HDsEMG placement, bipolar EMG electrodes for the right TA were positioned over a more proximal region of the muscle. All EMG signals were processed offline using notch filters at 50 and 100 Hz, followed by a fourth-order Butterworth band-pass filter (5–900 Hz).

### TMS settings

TMS was delivered over the left primary motor cortex using a monophasic magnetic stimulator (Magstim 200; Magstim Co., Whitland, UK) with a double-cone coil (outside diameter: 110 mm; Magstim Co., Whitland, UK). The induced current was oriented in the anterior-to-posterior direction. The coil was positioned over the optimal site (“hot spot”) for eliciting MEPs in the right TA. The position and direction of the coil were recorded with a navigation system (Brainsight; Rogue Research Inc., Montreal, QC, Canada) to ensure accuracy of coil positioning throughout the experiment. Resting motor threshold (rMT) was determined in the supine position and defined as the minimum intensity that evoked motor potentials with peak-to-peak amplitudes greater than 50 µV in at least 5 of 10 successive trials (Rossini et al. 2015) [48.4 ± 11.9 % maximum stimulator output (MSO) (mean ± SD)].

### tSCS settings

tSCS was delivered using a constant-current electrical stimulator (DS7A; Digitimer Ltd., United Kingdom) with a single monophasic square pulse (pulse width: 1 ms). The cathode electrode (50 × 50 mm) was placed over the thoracolumbar spine between the interspinous processes, and the anode electrode (100 × 75 mm) was placed on the abdomen above the umbilicus (Saito et al. 2019; Sasaki et al. 2020). Before data collection, the cathode was adjusted to determine the optimal stimulation location, producing the largest peak-to-peak responses across all recorded muscles. tSCS-evoked responses were tested at the T12–L1, L1– L2, and L2–L3 using the same stimulus intensity, and the optimal site was selected (T12–L1: n = 1, L1–L2: n = 9, L2–L3: n = 0). Electrodes were fixed with adhesive tape to prevent displacement during the experiments.

To determine stimulus intensity, recruitment curves were obtained for each participant by gradually increasing the tSCS intensity while recording responses from all muscles. To avoid ceiling effects in the evoked responses, the stimulus intensity was selected from the ascending part of the recruitment curve, ensuring that responses across muscles remained below saturation (Masugi et al. 2019; Sasaki et al. 2020), corresponding to approximately 1.3 × MT [58.6 ± 11.5 mA (mean ± SD)], where MT denotes the minimum stimulation intensity that consistently elicited reproducible spinally evoked motor potentials in the TA muscle.

To confirm that tSCS-evoked motor potentials were trans-synaptically mediated (Courtine et al. 2007), a paired-pulse protocol was applied before the main experiment (Steele et al. 2021) . Two tSCS pulses were delivered with a 50-ms interstimulus interval, repeated five times at the same stimulus intensity (1.3 MT). The second response was almost completely suppressed compared with the first response, and peak-to-peak amplitudes of the second responses were significantly smaller than those of the first responses for all recorded muscles (*P* < 0.05, Figure 2). This suppression is consistent with post-activation depression, and supports the interpretation that the responses were predominantly trans-synaptically mediated (Courtine et al. 2007; Dy et al. 2010; Masugi et al. 2016, 2017; Minassian et al. 2007; Sayenko et al. 2015b).

**Figure 2.**
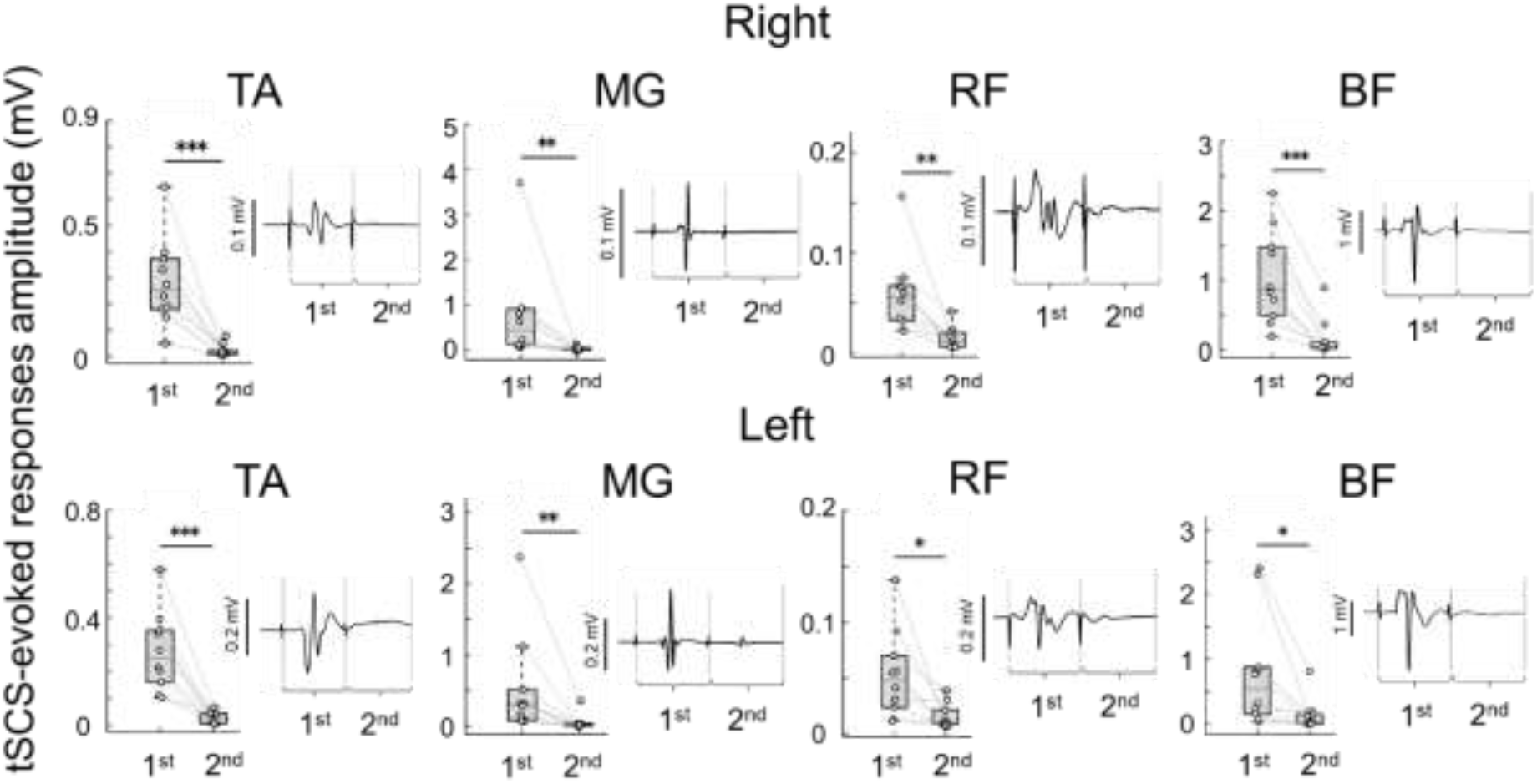
Group data for the first and second tSCS-evoked responses elicited at the ascending part of the recruitment curve. Data are shown for the tibialis anterior (TA), medial gastrocnemius (MG), rectus femoris (RF), and biceps femoris (BF) on both sides. * *P* < 0.05, ** *P* < 0.01, *** *P* < 0.001

### Experimental protocol

*Exp. 1* and *Exp. 2* were conducted on the same day with rest periods up to 15 min between sessions. The protocol for the *Exp. 1* is visualized in Figure 3. In *Exp. 1*, TMS-only stimulation at rMT and tSCS-only stimulation at 1.3 MT were each delivered five times (inter-trial interval: 10 sec). The onset latencies of the TMS-induced MEP and the spinally evoked motor potential in the right TA were visually determined from five trials using a LabChart (National Instruments, United States), and the mean latency across trials was calculated for each participant [MEP latency: 31.1 ± 1.9 ms; spinally evoked motor potentials latency: 16.8 ± 1.5 ms (mean ± SD)]. Central conduction time (CCT) was defined as the difference between the MEP latency and the spinally evoked motor potential latency. Interstimulus intervals (ISIs) between TMS and tSCS were defined for each participant based on the individual CCT. Eight ISI conditions were tested: CCT−10, CCT−5, CCT, CCT+5, CCT+10, CCT+15, CCT+20 and TMS+100 ms. For each ISI condition, baseline tSCS-only trials at 1.3 MT were obtained immediately before the combined stimulation trials. Combined stimulation consisted of TMS at 80% rMT combined with tSCS at 1.3 MT. ISI conditions were tested in randomized order. For both baseline (tSCS-only) and combined (TMS + tSCS) conditions, five trials were delivered per ISI with an intertrial interval of 10 seconds.

**Figure 3.**
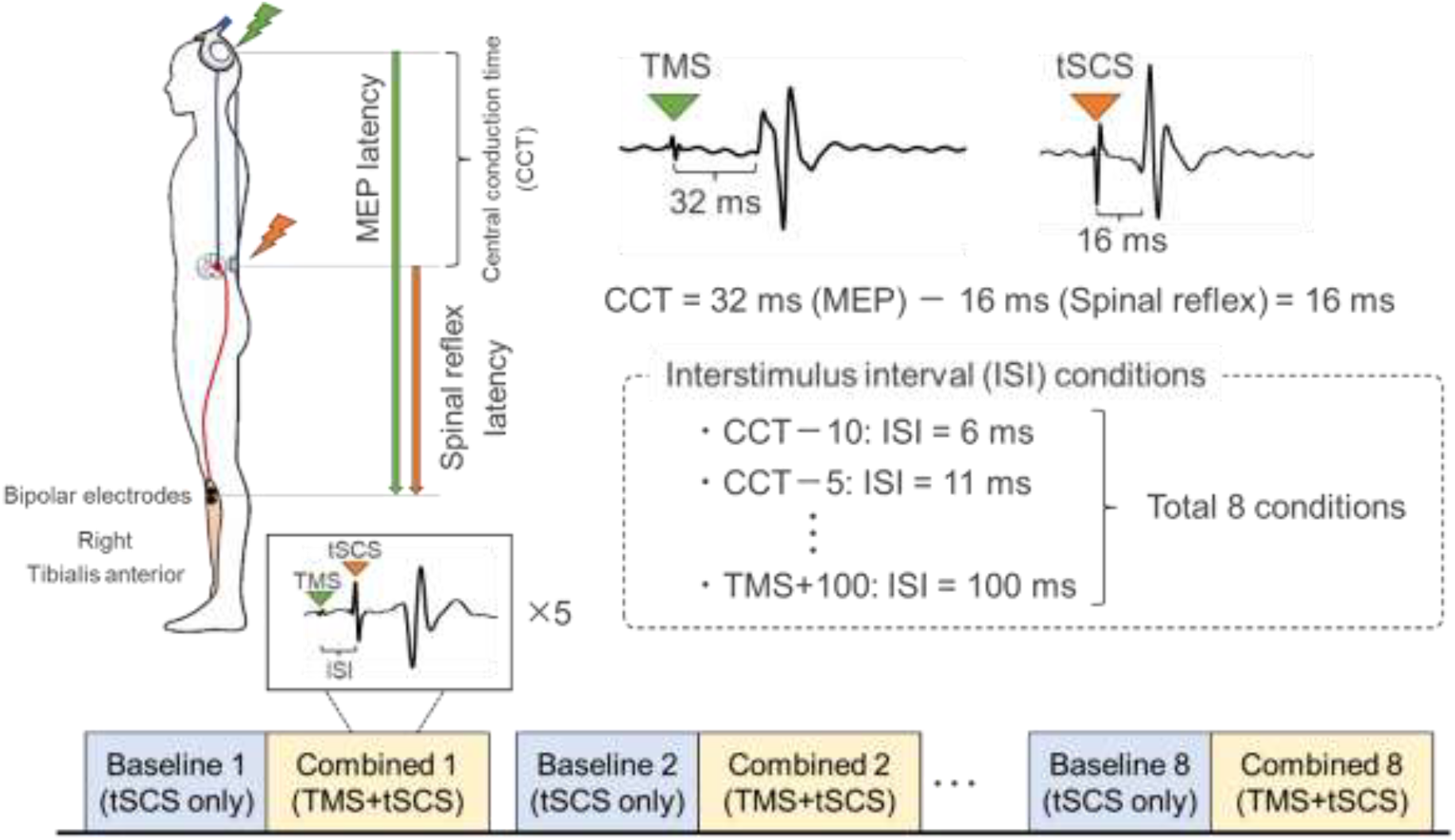
Experimental protocol for *Exp*.*1*. TMS-only stimulation and tSCS-only stimulation were delivered to calculate the latencies of the TMS-induced MEPs and spinally evoked motor potentials in right tibialis anterior, from which central conduction time (CCT) was calculated. Based on the CCT, eight interstimulus interval (ISI) conditions were determined. A positive ISI indicates that TMS was delivered prior to tSCS. For each ISI condition, baseline tSCS-only trials were delivered immediately before each combined TMS and tSCS condition. Each stimulation condition was repeated five times.

In *Exp. 2*, intramuscular activation of the right TA during the baseline (tSCS-only) and combined (TMS + tSCS) conditions was recorded using HDsEMG. The same ISI conditions as in *Exp. 1* were used for the baseline and combined conditions in *Exp. 2*. In addition to combined condition, single (TMS-only and tSCS-only) stimulation were each delivered five times (intertrial interval: 10 sec) to clarify and compare the intramuscular activation distribution evoked by each stimulus alone. During single stimulation sessions, to minimize the influence of response magnitude on spatial comparisons, stimulation intensities were adjusted so that the peak-to-peak amplitudes of the evoked responses in the right TA were comparable. Therefore, TMS at 1.2 rMT and tSCS at MT were performed in single stimulation sessions [TMS-induced MEP: 58.1 ± 14.3% MSO; spinally evoked motor potential: 46.8 ± 9.2 mA (mean ± SD)]. The single (TMS- and tSCS-only) stimulation sessions from one participant were not recorded due to the equipment malfunction, and analyses were conducted on the remaining nine participants.

### Data analysis

For *Exp. 1* and *Exp. 2*, peak-to-peak amplitudes were used to quantify TMS-induced MEPs and spinally evoked motor potentials in both bipolar EMG and HDsEMG recordings. These amplitudes were calculated using a custom-written MATLAB script (R2024a, The MathWorks Inc., Natick, MA, USA). For HDsEMG, peak-to-peak amplitudes were calculated separately for each of the 64 channels. Background EMG activity was quantified as the root mean square amplitude in the 50-ms window before the TMS or tSCS stimulus.

In *Exp*.*1*, the spinally evoked motor potentials amplitudes recorded with bipolar EMG during combined stimulation (TMS + tSCS) were normalized to the amplitudes measured during the immediately preceding baseline (tSCS-only) trials for each muscle across all ISI conditions.

In *Exp. 2*, HDsEMG was used to visualize intramuscular activation of the right TA during TMS-only, tSCS-only, and combined (TMS + tSCS) stimulation. For the TMS-only and tSCS-only conditions, heatmaps were generated from the channel-wise peak-to-peak amplitudes. To facilitate visualization of the intramuscular activation patterns, the channel exhibiting the largest response amplitude within the 64-channel HDsEMG grid was set to 1 for each participant, and all channels were normalized to this value. These participant-level normalized maps were then averaged across participants to generate group heatmaps for TMS-induced MEPs and spinally evoked motor potentials.

To characterize intramuscular facilitation during combined stimulation, facilitation maps were generated by subtracting the channel-wise spinally evoked motor potentials amplitudes obtained during baseline (tSCS-only) stimulation from those recorded during combined stimulation (TMS + tSCS) for each ISI condition. Group heatmaps were constructed by averaging the difference values across participants for each ISI condition.

To quantify the spatial distribution of the HDsEMG maps, the x- and y-coordinates of the center of gravity (CoG) were calculated across the 13×5 electrode grid for the TMS-only, tSCS-only, and facilitation map (combined − baseline). Specifically, the CoG coordinates were defined as:

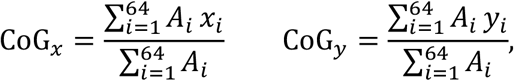

Where *A*_*i*_denotes the muscle activity amplitude recorded at the *i*-th electrode, and *x*_*i*_ and *y*_*i*_ represent the electrode positions of that channel along the mediolateral (column-wise) and proximodistal (row-wise) directions, respectively (Elswijk et al. 2008) . For the TMS-only and tSCS-only conditions, CoG was calculated from the channel amplitudes within each map. For the facilitation maps, CoG was calculated from the channel-wise difference values.

### Statistical analysis

#### Paired-pulse stimulation

In the paired-pulse protocol, normality of the EMG amplitudes was assessed using the Shapiro–Wilk test. Because the data were not normally distributed, the amplitudes of the first and second responses were compared for each lower-limb muscle using the Wilcoxon signed-rank test.

#### Background EMG

Background EMG activity was analyzed separately for each muscle using a linear mixed model (LMM). The dependent variable was the background EMG amplitude. Stimulation modality [i.e., baseline (tSCS-only) vs. combined (TMS + tSCS) stimulation], ISI conditions, and their interaction were included as fixed effects, and participant was included as a random effect.

#### spinally evoked motor potentials modulation in multiple lower-limb muscles at each ISI condition

To compare the spinally evoked motor potentials amplitude between baseline (tSCS-only) and combined (TMS + tSCS) conditions, an LMM was fitted separately for each muscle. The dependent variable was spinally evoked motor potentials amplitude, with stimulation modality, ISI condition, and their interaction included as fixed effects, and participant included as a random effect. When a significant interaction between stimulation modality and ISI condition was detected, post hoc pairwise comparisons between baseline (tSCS-only) and combined (TMS + tSCS) conditions were conducted separately for each ISI condition. *P*-values were adjusted using the Holm–Bonferroni correction.

#### Intramuscular activation patterns in the right TA muscle

To compare the spatial locations of TMS-induced MEPs, spinally evoked motor potentials and facilitation maps, the x- and y-coordinates of the CoG were analyzed separately at each ISI condition in which significant spinally evoked motor potentials facilitation were identified in *Exp. 1*. For each coordinate and ISI condition, an LMM was fitted with response type (TMS-induced MEP, spinally evoked motor potential, and facilitation) as a fixed effect and participant as a random effect. When a significant main effect of response type was detected, post hoc pairwise comparisons between response types were performed using Holm– Bonferroni correction. The significance level for all statistical tests was set at *P* < 0.05.

## Results

### Background EMG

As no significant main effects of stimulation modality, ISI condition, and their interaction on background EMG were observed in the LMM across all muscles (*P* > 0.05), there was no significant differences in background EMG between the baseline (tSCS-only) and combined (TMS + tSCS) conditions (Figure 4).

**Figure 4.**
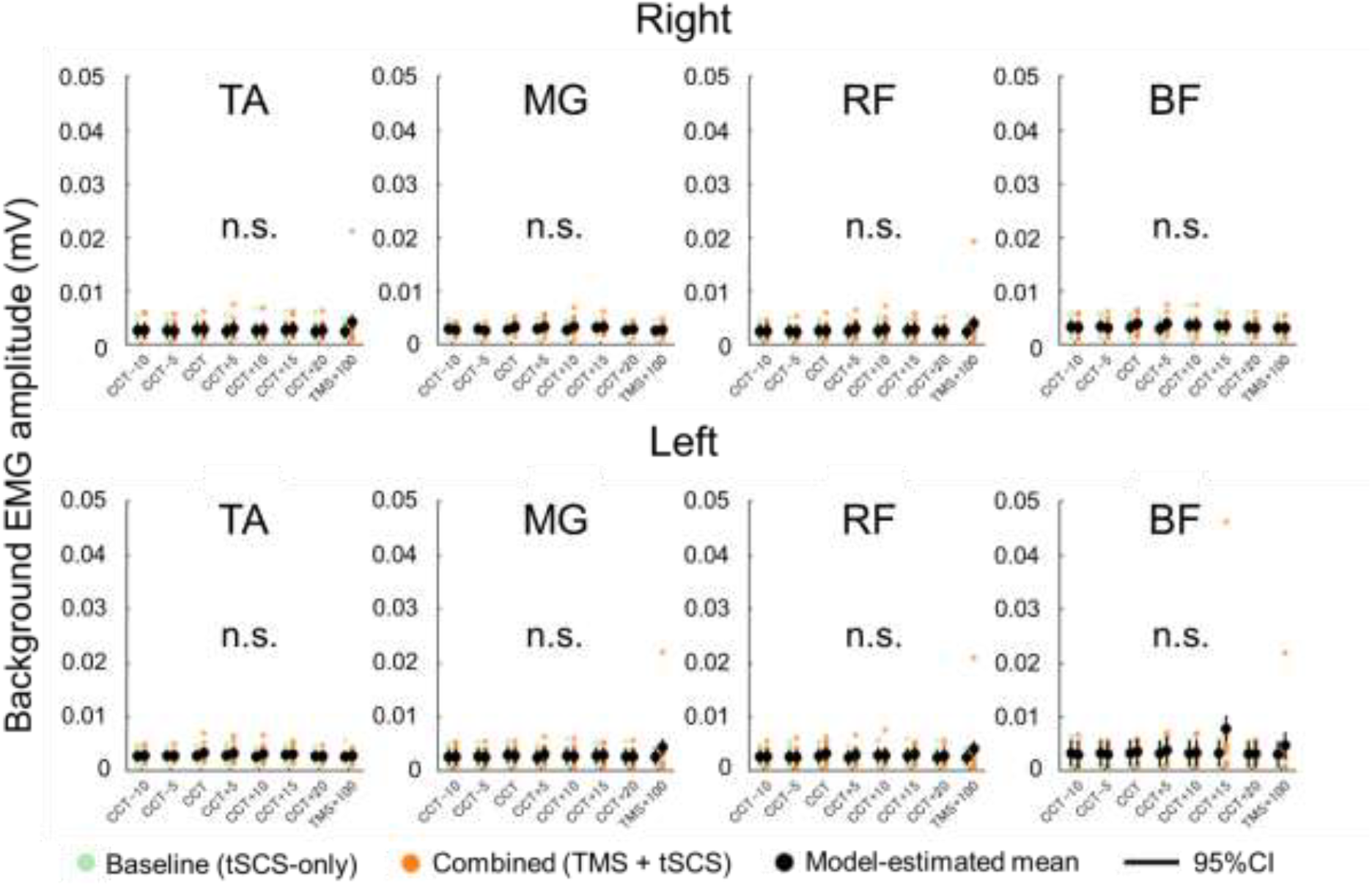
The results of linear mixed model (LMM) of background EMG activity in bilateral tibialis anterior (TA), medial gastrocnemius (MG), rectus femoris (RF) and biceps femoris (BF) during baseline (tSCS-only) and combined (TMS + tSCS) conditions. Green and orange data plots represent the mean values across five trials for each participant during baseline and combined conditions, respectively. Black plots represent the model-estimated means, and vertical black lines indicate the 95% confidence intervals (CI).

### Spinally evoked motor potentials modulation during combined TMS and tSCS stimulation

The LMM results of comparison of spinally evoked motor potentials amplitude between baseline (tSCS-only) and combined (TMS + tSCS) conditions are summarized in Table 1, and the descriptive statistics and post hoc pairwise comparisons with Holm-Bonferroni correction are shown in Table 2 and Figure 5A. Figure 5B shows representative spinally evoked motor potentials recorded from the right TA under each ISI condition, whereas Figure 5C shows representative responses from each lower-limb muscle at the CCT condition. Although the LMM showed no significant main effects of stimulation modality or ISI conditions, a significant interaction between stimulation modality and ISI condition was detected across all lower-limb muscles. Post hoc pairwise tests with Holm–Bonferroni correction revealed that spinally evoked motor potentials were significantly facilitated by TMS conditioning stimuli delivered at the CCT and longer ISI conditions in all lower-limb muscles (*P* < 0.05). In contrast, spinally evoked motor potentials in right BF were slightly suppressed at CCT−5 ms (98.0%, *P* = 0.008).

**Table 1.** The results of the LMM analyses examining the effects of stimulation modality, interstimulus interval (ISI) condition, and their interaction on lower-limb muscles. Since the interaction between stimulation modality and ISI condition was significant for all lower-limb muscles, post hoc analyses were performed for each muscle.

| Outcome | *P*-value | Outcome |  |
| --- | --- | --- | --- |
| Right |  | Left |  |
| TA |  | TA |  |
| Fixed effects |  | Fixed effects |  |
| Modality | 0.812 | Modality | 0.519 |
| ISI condition | 1 | ISI condition | 1 |
| Interaction |  | Interaction |  |
| Modality×ISI condition | < **0.001** | Modality×ISI condition | < **0.001** |
| MG |  | MG |  |
| Fixed effects |  | Fixed effects |  |
| Modality | 0.665 | Modality | 0.965 |
| ISI condition | 1 | ISI condition | 1 |
| Interaction |  | Interaction |  |
| Modality×ISI condition | < **0.001** | Modality×ISI condition | < **0.001** |
| RF |  | RF |  |
| Fixed effects |  | Fixed effects |  |
| Modality | 0.976 | Modality | 0.854 |
| ISI condition | 1 | ISI condition | 1 |
| Interaction |  | Interaction |  |
| Modality×ISI condition | < **0.001** | Modality×ISI condition | < **0.001** |
| BF |  | BF |  |
| Fixed effects |  | Fixed effects |  |
| Modality | 0.683 | Modality | 0.236 |
| ISI condition | 1 | ISI condition | 1 |
| Interaction |  | Interaction |  |
| Modality×ISI condition | < **0.001** | Modality×ISI condition | < **0.001** |

**Table 2.** Spinally evoked motor potential amplitudes across ISI conditions for the lower-limb muscles with tSCS intensity at 1.3 MT. Spinally evoked motor potential amplitudes and 95% confidence intervals (CI) are obtained from model-estimated means and expressed as percentages relative to the tSCS-only condition. *P*-values represent post hoc comparisons between the tSCS-only and combined TMS and tSCS stimulation at each ISI condition, corrected for multiple comparisons using the Holm–Bonferroni correction.

|  |  | ISI conditions |  |  |  |  |  |  |  |
| --- | --- | --- | --- | --- | --- | --- | --- | --- | --- |
|  |  | CCT<br>-10 | CCT-5 | CCT | CCT+5 | CCT<br>+10 | CCT<br>+15 | CCT<br>+20 | TMS<br>+100m<br>s |
| Right |  |  |  |  |  |  |  |  |  |
| TA |  |  |  |  |  |  |  |  |  |
|  | Amplitude (%) | 99.2 | 107.3 | 133.2 | 142.1 | 133.1 | 115.8 | 115.6 | 130.9 |
|  | 95% CI | (92.7-108.0) | (87.2-125.4) | (109.7-175.1) | (107.3-186.9) | (111.1-188.8) | (112.4-163.9) | (103.1-141.7) | (116.5-150.5) |
|  | *P*-value | 0.724 | 0.724 | **<0.001** | **<0.001** | **<0.001** | **<0.001** | **<0.001** | **<0.001** |
| MG |  |  |  |  |  |  |  |  |  |
|  | Amplitude (%) | 97.5 | 102.6 | 116.1 | 106.3 | 124.6 | 113.6 | 110.4 | 116.8 |
|  | 95% CI | (91.9-108.0) | (90.5-107.3) | (95.5-144.7) | (98.5-154.8) | (108.4-152.8) | (109.1-126.1) | (95.3-132.0) | (108.3-155.3) |
|  | *P*-value | 0.934 | 0.934 | 0.715 | **0.003** | **<0.001** | **<0.001** | **0.030** | **0.003** |
| RF |  |  |  |  |  |  |  |  |  |
|  | Amplitude (%) | 102.2 | 102.3 | 111.8 | 124.1 | 127.2 | 124.4 | 112.5 | 126.4 |
|  | 95% CI | (94.4-108.5) | (87.1-124.2) | (98.9-128.2) | (110.3-153.9) | (109.0-141.1) | (110.3-150.5) | (97.1-140.1) | (116.8-143.2) |
|  | *P*-value | 1 | 1 | **<0.001** | **<0.001** | **<0.001** | **<0.001** | **<0.001** | **<0.001** |
| BF |  |  |  |  |  |  |  |  |  |
|  | Amplitude (%) | 99.2 | 98.0 | 105.5 | 118.7 | 114.3 | 111.4 | 108.8 | 122.8 |
|  | 95% CI | (91.5-106.9) | (80.1-109.6) | (98.7-121.6) | (106.4-155.9) | (108.4-139.1) | (106.0-137.3) | (96.8-114.4) | (106.8-146.9) |
|  | *P*-value | 0.618 | **0.008** | **0.008** | **<0.001** | **<0.001** | **<0.001** | **0.028** | **<0.001** |
| Left |  |  |  |  |  |  |  |  |  |
| TA |  |  |  |  |  |  |  |  |  |
|  | Amplitude (%) | 100.2 | 98.9 | 111.7 | 106.1 | 109.4 | 113.4 | 103.6 | 124.8 |
|  | 95% CI | (90.6-116.5) | (86.9-110.8) | (96.9-134.6) | (99.6-173.2) | (101.4-136.8) | (99.6-118.0) | (101.7-127.7) | (105.4-142.4) |
|  | *P*-value | 0.642 | 0.642 | **<0.001** | **<0.001** | **<0.001** | **<0.001** | **<0.001** | **<0.001** |
| MG |  |  |  |  |  |  |  |  |  |
|  | Amplitude (%) | 101.4 | 97.5 | 107.2 | 109.1 | 108.4 | 104.7 | 107.2 | 119.6 |
|  | 95% CI | (94.4-123.1) | (88.4-106.1) | (97.7-147.2) | (95.7-149.1) | (93.0-136.7) | (92.6-118.1) | (95.8-129.5) | (107.0-138.7) |
|  | *P*-value | 1 | 1 | **0.041** | **<0.001** | 0.117 | 0.554 | 0.158 | **<0.001** |
| RF |  |  |  |  |  |  |  |  |  |
|  | Amplitude (%) | 102.0 | 96.4 | 100.2 | 114.2 | 108.9 | 109.5 | 109.3 | 106.4 |
|  | 95% CI | (96.6-106.8) | (83.0-101.8) | (90.3-120.1) | (100.1-149.2) | (97.6-126.2) | (99.1-150.0) | (97.2-125.7) | (95.2-140.1) |
|  | *P*-value | 0.828 | 0.574 | 0.597 | **<0.001** | 0.276 | 0.059 | **0.033** | 0.059 |
| BF |  |  |  |  |  |  |  |  |  |
|  | Amplitude (%) | 101.9 | 99.9 | 110.9 | 122.8 | 111.3 | 113.7 | 126.2 | 115.3 |
|  | 95% CI | (85.0-108.1) | (88.1-118.6) | (100.1-147.1) | (109.0-186.4) | (94.3-162.7) | (102.0-151.8) | (97.6-163.7) | (107.6-217.1) |
|  | *P*-value | 0.177 | 0.524 | **<0.001** | **<0.001** | 0.524 | **0.001** | **0.003** | **<0.001** |

**Figure 5.**
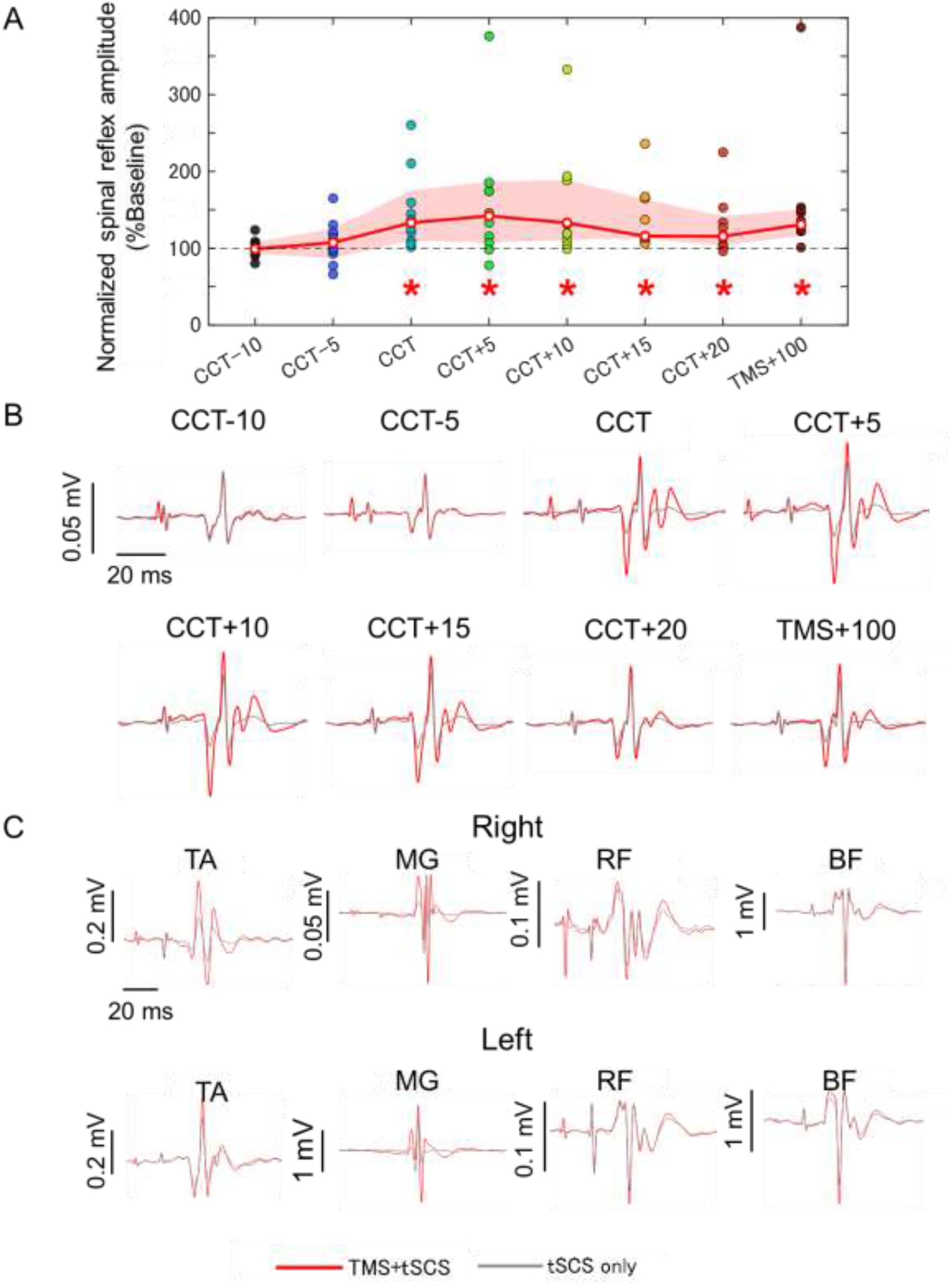
Spinally evoked motor potentials modulation by combined TMS and tSCS stimulation. (A) Normalized spinally evoked motor potentials amplitudes of the right TA at each ISI condition. White circles indicate the model-estimated means, and the red shaded band represents the 95% confidence interval derived from the linear mixed model. Asterisks indicate significant facilitation by combined TMS and tSCS stimulation (*P* < 0.05). (B) Representative spinally evoked motor potentials recorded from the right TA under each ISI condition. Red waveforms represent spinally evoked motor potentials during combined TMS and tSCS stimulation, and gray waveforms represent those during the baseline tSCS-only stimulation. (C) Representative spinally evoked motor potentials recorded from the bilateral TA, MG, RF and BF at the CCT condition.

### Intramuscular activation patterns of the right TA during TMS-only, tSCS-only and combined TMS + tSCS stimulation

Figure 6 shows the heatmaps and CoG locations of normalized TMS-induced MEPs, spinally evoked motor potentials, and facilitation maps at each ISI condition. For the y-coordinate of the CoG, the LMM revealed significant main effects of response type (TMS-induced MEP, spinally evoked motor potential, and facilitation) across all ISI conditions (*P* < 0.05), whereas no significant effects were observed for the x-coordinate (*P* > 0.1). Post hoc pairwise comparisons showed a significant difference in the y-coordinate of the CoG between TMS-induced MEPs and spinally evoked motor potentials (*P* = 0.003). The CoG of facilitation significantly differed from that of TMS-induced MEPs at the CCT+5, CCT+10, CCT+15, and TMS+100 ms (*P* < 0.05), and from that of the spinally evoked motor potentials at all ISI conditions (*P* < 0.001).

**Figure 6.**
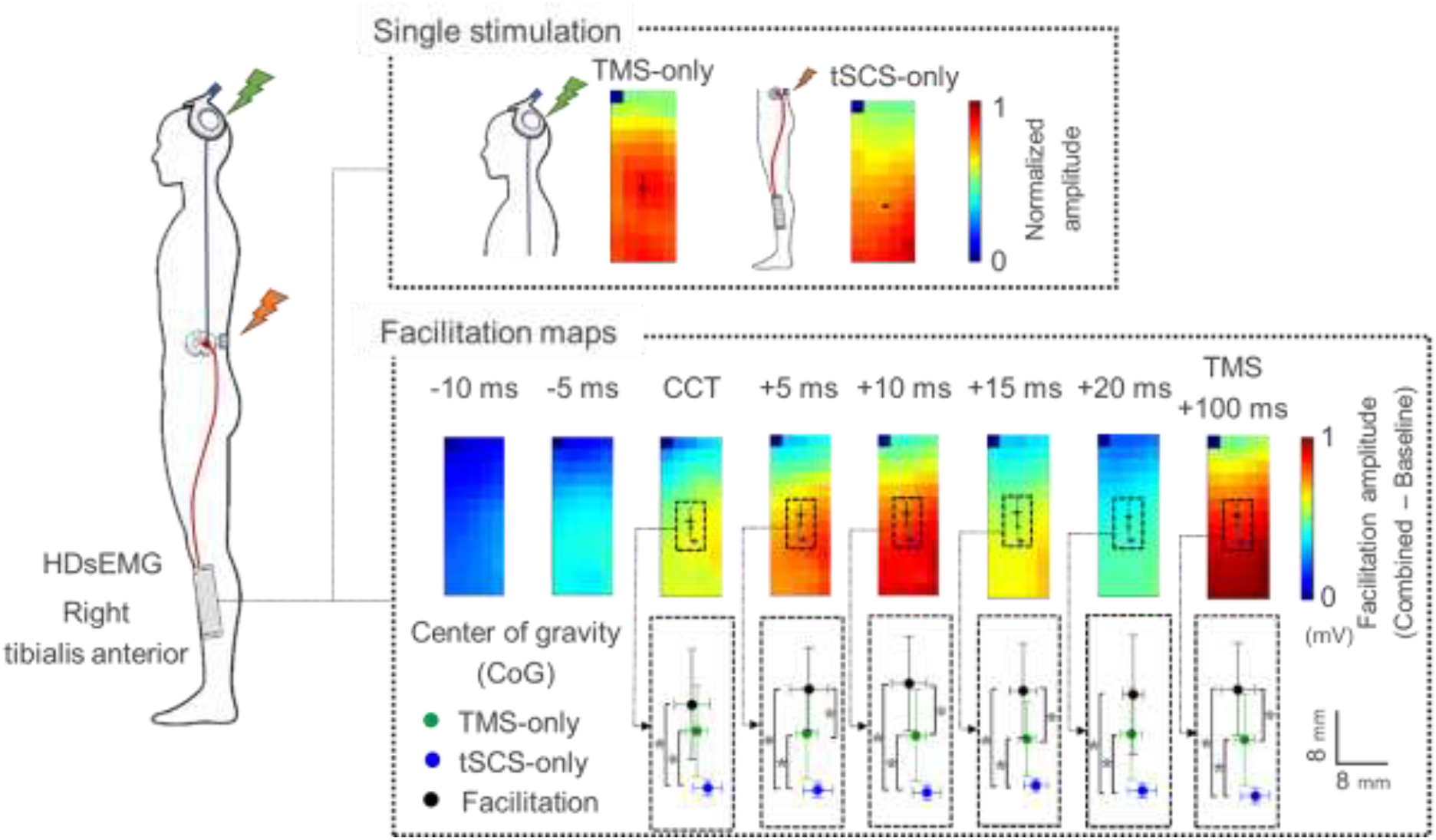
Heatmaps of intramuscular activation patterns in the right TA during TMS-only and tSCS-only stimulation, together with facilitation maps (combined − baseline) for each ISI condition. For the single-stimulation maps (TMS-only and tSCS-only), EMG amplitudes were normalized to the maximum channel value within each participant. Facilitation maps were generated on a channel-by-channel basis by subtracting the spinally evoked motor potentials amplitudes obtained during baseline (tSCS-only) stimulation from those obtained during combined (TMS + tSCS) stimulation at each ISI condition. Participant-level maps were then averaged across participants and displayed on a 13 × 5 grid. Colored circles indicate the CoG for TMS-only (green), tSCS-only (blue), and facilitation (black) maps. The color scale represents normalized amplitude for the single-stimulation maps and the facilitation amplitude (combined − baseline) for the facilitation maps with warmer colors indicating larger amplitudes.

## Discussion

The present study examined the spatiotemporal characteristics of descending modulation of multisegmental spinal network excitability in the lower limb by combining subthreshold TMS with thoracolumbar tSCS. In Experiment 1, subthreshold TMS facilitated spinally evoked motor potentials at CCT and longer ISI conditions in all recorded lower-limb muscles. Since background EMG activity did not differ significantly across any of the measured muscles between the baseline (tSCS-only) and the combined (TMS + tSCS) conditions, the observed modulation effects were likely attributable to the conditioning effect of TMS. In Experiment 2, HDsEMG revealed that the rostrocaudal position of the CoG of the MEP amplitude distribution differed from that of spinally evoked motor potentials. In addition, from CCT onward, the CoG of facilitation map was shifted proximally compared with those of TMS-induced MEP and tSCS-evoked motor potentials.

We hypothesized that spinally evoked motor potentials modulation is timing-dependent and could extend beyond the target muscle. Consistent with this, conditioning TMS facilitated tSCS-evoked motor potentials not only in the targeted TA but also in non-target, including contralateral, muscles. To our knowledge, this study is the first to demonstrate that, under the present stimulation configuration, the descending volley elicited by TMS influenced spinal excitability across multiple motor pools projecting to the lower-limb muscles.

Several mechanisms may contribute to these multi-muscle and bilateral effects. Although corticospinal projections are predominantly contralateral, ipsilateral influences can arise via uncrossed corticospinal fibers and via polysynaptic spinal and brainstem relays that can be recruited by a TMS-induced cortical volley (Jankowska and Edgley 2006; Nielsen et al. 2007; Ziemann et al. 1999). In addition, TMS over M1 may engage cortico-reticular pathways and thereby influence reticulospinal output, which projects bilaterally and diffusely to spinal circuits (Peterson et al. 1975; Sakai et al. 2009). Finally, spinal commissural interneuronal networks can distribute excitation across side and adjacent spinal segments (Matsuyama et al. 2006). Through these pathways, a subthreshold conditioning volley could increase the net excitability of multiple motoneuron pools and thus increase the spinally evoked motor potentials evoked by a given afferent input (Sayenko et al. 2018; Taccola et al. 2025) .

From a functional perspective, such broad descending modulation may be relevant to the neural control during upright stance and locomotion, which requires coordinated bilateral and multi-joint actions rather than isolated single-muscle control. Reticulospinal output increases during anticipatory postural adjustments in standing (Chiou et al. 2024) and voluntary gait modifications, such as obstacle avoidance during walking (Prentice and Drew 2001), and commissural interneurons, which receive substantial input from the reticulospinal tract, regulate left–right alternation during locomotion (Maxwell and Soteropoulos 2020). Accordingly, a descending volley that biases excitability across multiple motoneuron pools could provide a physiological substrate for rapidly adjusting reflex responsiveness across the lower-limb musculature in behaviorally relevant contexts.

At the same time, methodological factors may also have contributed to the broad facilitation observed here. Selective activation of a single lower-limb muscle is difficult with TMS, particularly when using a double-cone coil, because the induced electric field can spread to neighboring representations (Kesar et al. 2018). Therefore, the conditioning pulse may have engaged corticospinal outputs related to multiple lower-limb muscles, which would also be expected to promote facilitation across the recorded muscles.

Our hypothesis that TMS-conditioned modulation of spinally evoked motor potentials is timing-dependent was also validated. Consistent with previous studies, spinally evoked motor potentials were facilitated from CCT through CCT+20 ms (Andrews et al. 2020; Rossi et al. 2025; Roy et al. 2014; Sayenko et al. 2018). Importantly, facilitation was still evident even at the considerably longer ISI of TMS+100 ms. Whereas previous studies defined stimulus interval based on the timing difference between peripheral nerve stimulation and TMS (i.e. C-T intervals) (Andrews et al. 2020; Lopez et al. 2020; Niemann et al. 2018; Rossi et al. 2025), the present study determined interstimulus intervals using the latency difference between TMS-induced MEPs and spinally evoked motor potentials. Because differences in stimulation site and afferent pathways introduce latency shifts of several milliseconds (up to ∼10 ms) between the H-reflex and tSCS responses (Minassian et al. 2007), the observed facilitation at and beyond CCT is broadly consistent with prior reports showing H-reflex facilitation occurring several milliseconds after peripheral nerve stimulation (Andrews et al. 2020; Niemann et al. 2018). At CCT and CCT+5 ms conditions, facilitation likely reflects temporal summation between fast descending input evoked by TMS and Ia afferent input evoked by tSCS at the level of spinal motoneuron pool (Nielsen and Petersen 1995; Pierrot-Deseilligny and Burke 2005; Rothwell et al. 1991). At CCT+10, +15, and +20 ms, slower polysynaptic corticospinal inputs may increase motoneuronal excitability before the arrival of the tSCS-evoked afferent volley, thereby, facilitating the spinal responses (Andrews et al. 2020; Rossi et al. 2025). The mechanism underlying this delayed facilitation at TMS+100 ms remains unclear. One possible explanation is that TMS indirectly engages brainstem-mediated descending influences, including reticular structures, which may modulate descending drive to lumbar motoneurons (Heckman et al. 2008) and activate reticulospinal pathways projecting bilaterally to these motoneurons (Akalu et al. 2023; Fisher et al. 2021; Peterson et al. 1975). This may shift motoneurons toward a state in which persistent inward currents are more readily expressed (Heckman et al. 2008), thereby amplifying subsequent Ia afferent input elicited by tSCS and enhancing spinally evoked motor potentials.

Consistent with our hypothesis, intramuscular activation patterns differed significantly between TMS-induced MEPs and spinally evoked motor potentials. Although both heatmaps showed prominent activation in the distal region of the right TA, TMS-induced MEPs exhibited relatively greater activation in the central region of the muscle compared to spinally evoked motor potentials. Accordingly, the CoG of TMS-induced MEPs was located more proximally than that of spinally evoked motor potentials. Recent studies suggest that the sets of motoneurons innervating the proximal and distal regions of the TA muscle receive distinct synaptic inputs (Weinman et al. 2026). Moreover, CoG shifts might reflect changes in motor unit recruitment within the same muscle (Elswijk et al. 2008; Falla and Gallina 2020; Farina et al. 2008). The findings in the present study suggest that, although both descending and Ia afferent inputs excite motoneurons innervating the distal region of the TA muscle, they may preferentially recruit distinct subsets of motoneurons innervating different regions within the same muscle, as reflected by the intramuscular spatial distribution of activation.

To characterize the intramuscular patterns of facilitation of spinally evoked motor potentials induced by subthreshold TMS conditioning, facilitation maps (combined – baseline) were generated for ISI conditions from CCT onward, where significant facilitation of spinally evoked motor potentials was observed in *Exp*.*1*, and the corresponding CoG was calculated. The CoG of the facilitation maps was located more proximally than that of the spinally evoked motor potentials and, in most ISI conditions, also more proximally than that of TMS-induced MEPs. This proximal shift indicates that facilitation during combined stimulation was spatially non-uniform relative to the activation pattern evoked by tSCS alone.

One possible explanation for the proximal bias of facilitation is that a subset of motoneurons may not have been fully recruited by afferent input alone in more proximal regions, and that subthreshold TMS may have transiently increased the excitability of motoneurons innervating these proximal regions. The temporal interaction between TMS-induced descending inputs and afferent input may therefore preferentially enhance activation of proximal motoneuron populations during combined stimulation.

## Conclusion

Descending volleys evoked by TMS facilitated spinally evoked motor potentials when their arrival at the motoneuron pool was temporally aligned, consistent with convergent integration of descending and afferent inputs at the level of spinal interneuronal networks and motoneurons. This facilitatory effect was not confined to the target muscle but extended across multiple lower-limb muscles, indicating that corticospinal inputs can modulate excitability across multiple segments in a distributed manner, likely via shared premotor interneuronal circuitry. Within the TA, TMS-induced MEPs and spinally evoked motor potentials exhibited broadly similar intramuscular activation patterns but differed in their CoG, suggesting distinct spatial weighting of motor unit recruitment under descending versus afferent drive. Notably, facilitation during combined stimulation was preferentially expressed in proximal regions of the muscle, indicating that the interaction between descending and afferent inputs is spatially non-uniform and may reflect region-specific differences in motoneuron recruitment thresholds, synaptic gain, or saturation of afferent-driven activation. Together, these findings support a model in which temporally aligned corticospinal-afferent coupling produces synergistic amplification of motor output through non-linear integration within spinal sensorimotor circuits.

